# Atypical domain communication and domain functions of a Hsp110 chaperone

**DOI:** 10.1101/220798

**Authors:** Vignesh Kumar, Joshua Jebakumar Peter, Amin Sagar, Arjun Ray, Ashish, koyeli mapa

## Abstract

Hsp110s are well recognized nucleotide exchange factors (NEFs) of Hsp70s, in addition they are implicated in various aspects of cellular proteostasis as discrete chaperones with yet enigmatic molecular mechanism. Stark similarity in domain organization and structure between Hsp110s and Hsp70s, is easily discernible although the nature of domain communication and domain functions of Hsp110s are still puzzling. Here, we report atypical domain communication of yeast Hsp110, Sse1 using single molecule FRET, small angle X-ray scattering measurements (SAXS) and Molecular Dynamic simulations. Our data show that Sse1 lacks typical domain movements as exhibited by Hsp70s, albeit it undergoes unique structural alteration upon nucleotide and substrate binding. Hsp70-like domain-movements can be artificially salvaged in chimeric constructs of Hsp110-Hsp70 although such salvaging proves detrimental for the NEF activity of the protein. Furthermore, we show that substrate binding domain (SBD) of Hsp110, chaperones self, as well as foreign nucleotide binding domains (NBD). Interestingly, the substrate binding specificity of Hsp110 is largely determined by its NBD rather than SBD, the latter being the foremost substrate binding region for Hsp70s.

## Introduction

Heat Shock protein 70s (Hsp70s) are extremely conserved and ubiquitously present group of molecular chaperones. These proteins are involved in multitude of cellular processes and are assisted and regulated by two groups of co-chaperones to accomplish their functions in physiological time scales: J-domain proteins or Hsp40s and nucleotide exchange factors (NEFs) (Hartl, Bracher et al., 2011, Hartl & Hayer-Hartl, 2009, Kim, Hipp et al., 2013). While Hsp40s accelerate the weak ATPase rate of Hsp70s, NEFs accelerate the process of exchange of hydrolysed ADP with fresh ATP molecules, facilitating the initiation of a new chaperone cycle. Hsp110 proteins have been shown to act as NEFs for Hsp70s and are exclusively found in eukaryotes along with at least three other groups of structurally dissimilar NEFs, the GrpE homologs found in sub-cellular compartments of endo-symbiotic origin, HspBP1 proteins (like Fes1 in yeast) and Bag domain NEFs (Bracher & Verghese, 2015). Unlike these three types of NEFs, Hsp110s possess extremely similar domain organization and molecular architecture with the Hsp70s. The structural intricacies along with molecular mechanism of NEF activity of Hsp110s were nicely captured by a series of studies on yeast Hsp110, Sse1 (Andreasson, Fiaux et al., 2008, Bracher & Verghese, 2015, Liu & Hendrickson, 2007, Polier, Dragovic et al., 2008, Schuermann, Jiang et al., 2008). In these studies, Sse1 structure was solved in the ATP or ATP-analogue-bound states either in isolation or in complex with various Hsp70 partners (Liu & Hendrickson, 2007, Polier et al., 2008, Schuermann et al., 2008). In the ATP or ATP-analogue bound states, Sse1 was found in a lid-open structure with extensive contacts between the two domains; nucleotide binding domain (NBD) and substrate/peptide binding domain (SBD/PBD) (Liu & Hendrickson, 2007, Polier et al., 2008, Schuermann et al., 2008). On the other hand, the two-domain structures of Hsp70s were rare in literature till recent years probably due to dynamic nature of Hsp70 domains. The ATP-bound two-domain structure came out only recently (Qi, Sarbeng et al., 2013), previous to this structures of ATP-bound states could be solved using hydrolysis deficient mutants of Hsp70s or using non-hydrolysable ATP analogues plausibly due to intrinsic ATP hydrolysis by Hsp70s over the course of the experiment (Kityk, Kopp et al., 2012, Zhuravleva, Clerico et al., 2012). The structure of ADP-bound-states of Hsp70s (without bound substrate) are extremely rare in literature. Single molecule FRET studies on different Hsp70s could capture the conformational heterogeneity ADP-bound states of the chaperone, plausibly due to dynamicity in the ADP state, explaining the difficulty in getting crystals in ADP-state of the Hsp70s. What is clear from all such studies is that Hsp70s undergo prominent domain movements resulting in allosteric regulation of domain functions, crucial for its chaperone function. In contrast, structural insights about Sse1 or other members of Hsp110 chaperones in alternate functional states (e.g. in ADP or substrate-bound states), are so far lacking in literature except a study where the authors had employed HD-X coupled to Mass Spectrometry on Sse1 in both ADP and ATP- states (Andreasson et al., 2008). It was demonstrated that the signature of tryptic fragments of Sse1 in ATP and ADP-bound states were nearly identical leading to the conclusion that Sse1 lacks ATP-hydrolysis driven domain allostery (Andreasson et al., 2008). In the context of other cellular functions attributed to Hsp110s (Hrizo, Gusarova et al., 2007, Liu, Morano et al., 1999, Makhnevych, Wong et al., 2012, Mandal, Gibney et al., 2010, Sadlish, Rampelt et al., 2008), especially its role in prevention of aggregation and degradation of misfolded proteins (Kuo, Ren et al., 2013, Zhang, Binari et al., 2010), it is yet unclear how Hsp110s bind and release such misfolded/aggregated proteins in absence of a canonical allosteric regulation.

Here, by employing single molecule FRET measurements (sm-FRET) in solution, we report that Hsp70-like canonical domain movements are absent in Hsp110, although we are able to capture unique changes in conformation in the ADP and ATP-states of the protein using SAXS measurements and MD simulation. SAXS models have given us new insights into the substrate-binding induced structural alterations of this unique group of chaperones. Interestingly, in chimeric constructs of Hsp110-Hsp70, the canonical Hsp70-type domain movements are regained, however such domain movements abolish the NEF activity albeit keeping the chaperoning activity of the chimeric proteins intact, during proteotoxic stresses. We further show that not only the domain communication but also the functions of Hsp110 domains have significantly diverged from their Hsp70- counterparts. While the substrate binding domains (SBD) of Hsp110 harbor strong chaperoning activity on nucleotide binding domains (NBD) thereby playing a crucial role in stabilizing the protein, NBDs play an important role in substrate binding and determining the binding specificity of Hsp110s. The role of NBD in substrate binding, especially in determining the binding specificity is unusual for Hsp70s. In summary, here we report atypical domain movements and domain functions of Hsp110 proteins which are significant deviation from canonical mechanism of domain allostery of Hsp70s as well as its domain functions.

## Results

### Yeast Hsp110, Sse1, lacks prominent nucleotide dependent canonical domain movements like Hsp70s

The first high resolution structure of a member belonging to Hsp110 chaperones (yeast Hsp110, Sse1) came into light about a decade ago (Liu & Hendrickson, 2007). Thereafter, many other groups have solved the structure of Sse1 in complex with its various Hsp70 partners (Andreasson, Rampelt et al., 2010, Polier et al., 2008, Schuermann et al., 2008). Notably, all such high resolution structures of Sse1, either in isolation or as complex with its Hsp70 partners have been solved in the ATP or ATP-analogue-bound states, although the nature of ATP-hydrolysed state of Hsp110s remained enigmatic so far. A study using HDX-coupled to mass spectrometry by Andreasson *et al* had shown that the deuterium exchange rates of tryptic fragments of the Sse1 is similar in the ADP and the ATP-bound states of the protein (Andreasson et al., 2008). From this finding, the authors concluded that Sse1 lacks typical Hsp70-like allosteric communication between its domains driven by bound nucleotides. In this context, it is important to reiterate that previous observations from our group as well as from other groups with multiple Hsp70s in single molecule resolution have revealed that the ADP bound states of Hs70s are inherently heterogeneous compared to the ATP-bound states (Banerjee, Jayaraj et al., 2016, Mapa, Sikor et al., 2010, Marcinowski, Holler et al., 2011). This finding made us curious to check whether such conformational heterogeneity of the ADP-bound state is conserved in case of Hsp110s as well which possibly will remain masked during ensemble measurements like HDX-MS. Thus, to monitor conformational alterations in single molecule resolution, we subjected yeast Hsp110, Sse1, for single molecule FRET measurements to detect any large alteration in inter-domain or intra-domain distances which might result from allosteric communication between or within domains. To be able to subject the protein for smFRET measurements, we need to label the protein with donor-acceptor fluorophore pairs at specific sites. For that purpose, we usually substitute non-conserved surface exposed residues to cysteines to be able to label with maleimide-reactive fluorophores. In order to label Sse1, we substituted the native cysteine residues to alanine and as shown before, the cysteine substituted Sse1 protein is functional (Andreasson et al., 2008). On the cysteine-less version of Sse1, we engineered the cysteine pairs at desired location for fluorophore labelling. We made three such pairs to monitor the changes in distance due to possible conformational alterations: 1. between NBD and SBD (E319C-D412C), 2. between the lid and base of SBD (D412C-D600C) and 3. between the SBD-lid and NBD (K45C-K600C) (Figure 1A). We took single molecule FRET measurements of donor-acceptor labelled Sse1 proteins in solution in extremely low concentration (~50pM) to achieve the single molecule resolution as described before (Mapa, Tiwari et al., 2012).

**Figure 1:**
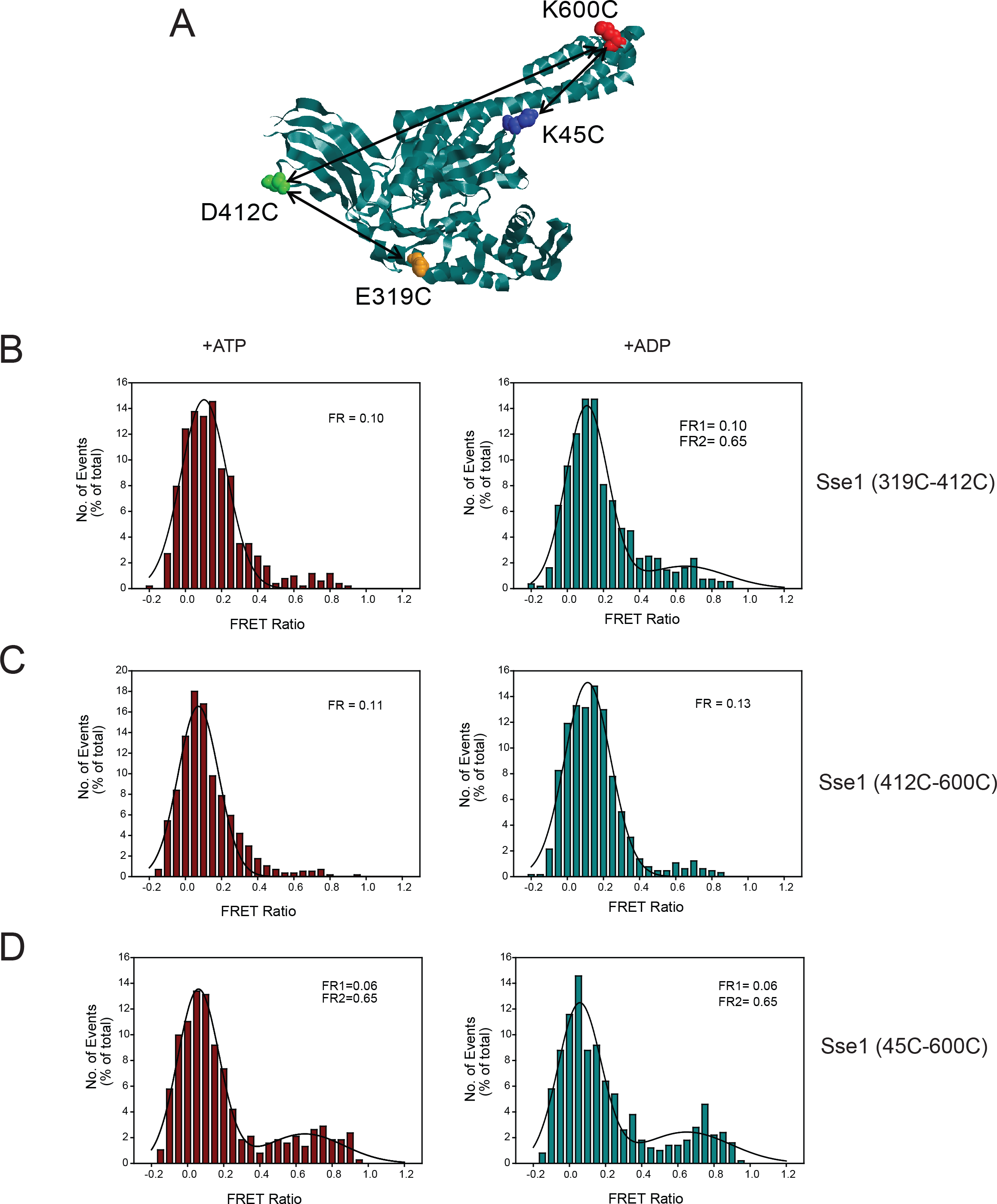
Domain movements of Sse1 probed by single molecule FRET experiments. (A). Structure of Sse1 (PDB ID: 2QXL) in cartoon representation showing residues substituted with cysteine for donor-acceptor label incorporation. Arrows indicate FRET pairs used to monitor domain movements. (B) – (D) smFRET ratio histograms showing population of Sse1 in various conformational states during ATP (Left panel) and ADP (Right panel) bound condition.(B) smFRET ratio histograms of Sse1 (319C-412C) and (D) Sse1 (45C, 600C) containing FRET pairs between PBD and NBD show a low FRET ratio population in both ATP and ADP bound condition indicating lack of domain docking-undocking movements as observed in canonical Hsp70s. (C) smFRET measurements on Sse1 (412C–600C) with FRET pairs between β-sheet subdomain and α helical lid of PBD show significant low FRET population during ATP and ADP bound states indicating a lid open conformation of the PBD in both the states.

To check for nucleotide-dependent alterations in inter-domain distance as a result of allosteric communication between NBD and SBD, we subjected Sse1 (319C-412C) for sm-FRET measurements. In the ADP-bound state, we observed a heterogeneous distribution of molecules. Majority of molecules populated at low FRET Ratio region with peak at FR, 0.1. Additionally, a small population of molecules were also present at high FR region (FR = 0.65) (Figure 1B, right panel). Surprisingly, the FR distribution of molecules remained essentially the same in the ATP bound state as well (Figure 1B, left panel). This finding is in stark contrast to Hsp70s where it was observed that SBD docks on to NBD in the ATP-state of the Hsp70s resulting in high FR molecules and in the ADP-state, SBD separates from NBD, thus shifting the population of molecules to low FR region (Mapa et al., 2010, Marcinowski et al., 2011). This change in inter-domain distance was nicely captured by sm-FRET measurements for at least two members of Hsp70 family where the shift of majority of molecules from high (ATP state) to low (ADP state) FRET ratio/efficiency peak were clearly discernible (Mapa et al., 2010, Marcinowski et al., 2011). In contrast, in case of Sse1 in spite of ATP-hydrolysis, we do not see any alteration of FR of molecules indicating no significant alteration in inter-domain distance in case of Hsp110s.

Next, we went ahead with the SBD-lid-base mutant (D412C-D600C). With this mutant, we observed a single prominent peak at low FR region (FR = 0.1) in the ATP state indicating an open lid conformation of SBD. This was expected as the crystal structures had reported the widely open structure of the Sse1 lid in the ATP-state. The ADP-state also was nearly identical in FR distribution indicating absence of prominent intra-domain conformational change within SBD due to lid movements post ATP-hydrolysis (Figure 1C).

While generating mutants for sm-FRET studies, we reasoned that if the lid of the SBD (which is wide open in ATP state and is positioned close to NBD) is dynamic like the lid of Hsp70s, the distance between the lid and NBD will change according to bound nucleotides. So a third reporter was constructed to report for any change in distance between the SBD-lid and NBD (K45C-K600C). This mutant revealed prominent bimodal distribution with FR peaks at 0.1 and 0.65 FR region indicating heterogeneity in SBD-lid-NBD distance. This conformational distribution remained unchanged with change of nucleotides and were essentially same in both ATP and ADP states (Figure 1D). This data indicates that indeed the SBD-lid is wide open and comes near NBD although the lid might be dynamic irrespective of the nucleotide state of the chaperone. The exact nature of such lid dynamicity and its implication on chaperoning activity of Sse1 needs further exploration. Altogether, our sm-FRET data with Sse1 shows that, the domain communication in case of Hs110s is unique and does not recapitulate the nucleotide dependent domain communication of Hsp70s. As we could not detect any noticeable ATP-hydrolysis dependent changes in inter-domain or intra-domain (SBD) distances as already described for multiple Hsp70s by single molecule FRET measurements, we reasoned that the nucleotide induced conformational changes in Hsp110, if any, might be too subtle to capture by this technique or are significantly different from the ones exhibited by the Hsp70 partners.

### Small Angle X-ray scattering (SAXS) measurements revealed distinctive nucleotide-dependent conformational changes of Hsp110

To detect any non-canonical conformational alteration during chaperone cycle of Hsp110s, we subjected Sse1 for SAXS measurements to determine large scale changes in domain orientation. To investigate the effect of nucleotides on the solution conformation of Sse1, we acquired and analyzed SAXS data in nucleotide bound states under saturating concentrations of ATP and ADP (please see methods section) as well as in the apo state of the molecule. The SAXS intensity profiles showed no upward or downward profile at low angles indicating that the samples were free of aggregation and there was no inter-particle interference in any of the conditions (Figure 2A). Linear profile of the Guinier analysis confirmed mono-dispersity of the scattering species profile (Figure 2A inset). Slope of the linear fits provided radius of gyration (Rg) of Apo, ATP and ADP enriched states to be 3.71±0.14, 3.73±0.47 and 3.77±0.38 nm, respectively corroborating the FRET results of no major conformational changes depending on the nature of the bound nucleotide. Computed P(r) curves for Apo, ATP and ADP enriched states showed a maximum linear dimension (Dmax) and Rg values of 13, 12 and 12.2, and 4.0, 3.8 and 3.8 nm, respectively (Figure 2B). Normalized Kratky plots for all three states had a maxima close of 1.73 supporting that the proteins have globular folded profile (Figure 2B inset). The shape parameters indicated no large scale shape change in the Sse1 protein as a function of bound nucleotides.

**Figure 2:**
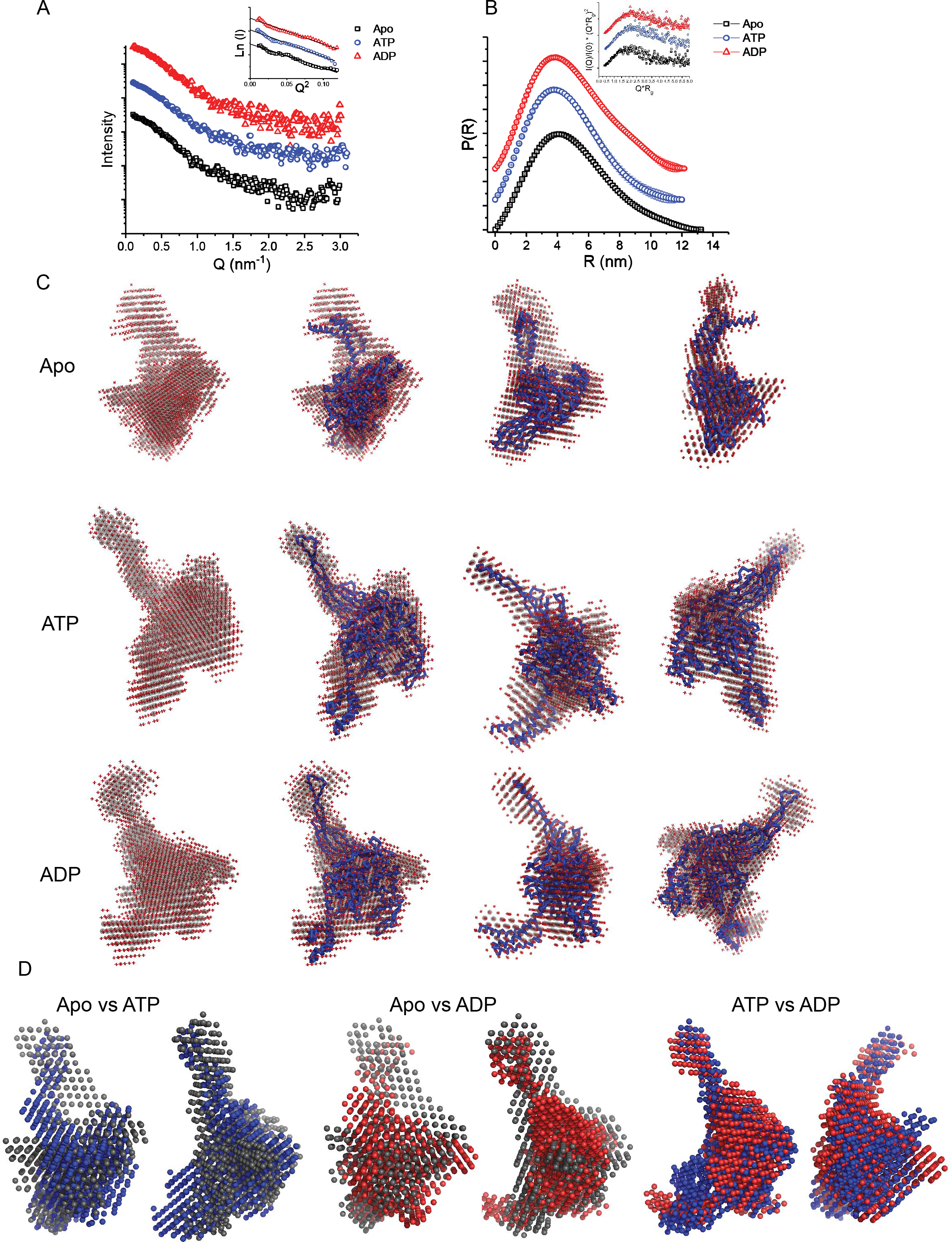
SAXS profile of Apo and ATP/ADP bound forms of Sse1. (A) SAXS intensity profiles of Sse1 in Apo form (Apo-Sse1) and bound to ATP (ATP-Sse1) or ADP (ADP-Sse1) are plotted as I(Q) (log scale) vs. Q (linear scale) with the Guinier Plots (ln I(Q) vs. Q) in the inset. The black solid lines in the Guinier plot show the linear fits. (B) The distance distribution curves (P(R) vs. R) derived by Indirect Fourier transformation of the intensity profiles with the normalized Kratky plots (I(Q)/I(0)*(Q*s)^2^ vs. Q*Rg) in the inset. (C) Starting from top, the first column shows the damfilt (Dark grey) and damaver (Red) models derived after the averaging of 10 dummy residue models for Apo-Sse1, ATP-Sse1 and ADP-Sse1, respectively. The following three columns show three orthogonal views of the SAXS derived models of Apo-Sse1 overlaid with the crystal structure of ATP bound form (PDB ID: 2QXL) using inertial axis alignment in the top row and the SAXS derived models of ATP-Sse1 and ADP-Sse1 overlaid with the ADP-BeFX bound crystal structures in the middle and bottom rows, respectively (PDB ID: 3C7N). (D) Starting from left, two orthogonal views each of the overlay of the SAXS models of Apo-Sse1 (Dark grey) with ATP-Sse1 (Blue), ApoSse1 with ADP-Sse1 (Red) and ATP-Sse1 with ADP-Sse1.

To visualize the global shape of the Sse1 protein with or without bound nucleotides, SAXS data profile was used to restore the scattering shape of the molecule under Apo and ADP or ATP bound state. Chain-like ensemble calculations were done to restore ten different models for each state and were averaged after alignment of their inertial axes. The values of average Normalized Spatial Discrepancy with the corresponding standard deviations between the 10 models were 0.55 ± 0.007, 0.76 ± 0.009 and 0.86 ± 0.007 for Apo, ATP-Sse1 and ADP-Sse1, respectively, indicating that the modelling procedure was stable and reproducible. Upon superimposition, it was clear that the model for Apo state was comparable with crystal structure of Sse1 in the ATP-state (PDB ID: 2QXL) with NSD of 1.3 (Figure 2C, top row) (Liu & Hendrickson, 2007). On the other hand, the ATP or ADP bound states were more similar to Sse1 crystal structure in ADP-BeFX (non-hydrolysable ATP analogue) in complex with bovine Hsc70 NBD (PDB ID: 3C7N) with NSD of 1.2 in contrast to ATP bound structure (PDB ID: 2QXL) with NSD of 2.5 (Figure 2C, second and third row) (Schuermann et al., 2008). Altogether, comparison of the SAXS based models confirmed that Sse1 protein remains as monomer and its shape largely remains unchanged regardless of the bound nucleotide status. The SAXS data very nicely correlated with sm-FRET data with three different FRET-based reporters which were used to report for NBD-SBD interaction or SBD-lid opening-closing movements, due to change in bound nucleotides. Although, there are no significant conformational changes in Sse1 due to ATP hydrolysis as seen in Hsp70s, analysis of SAXS - based models revealed that the nucleotide bound state significantly deviates from the Apo state. Importantly, comparison of nucleotide-bound conformation with the apo-state showed that nucleotide binding induces extended conformation of β-sheet region in the base of SBD. This data show that nucleotide binding imparts atypical conformational changes which might be characteristic of Sse1 or Hsp110s in general, the significance of such conformational changes for chaperoning function of the protein although remain elusive.

Altogether, sm-FRET spectroscopy and SAXS measurements collectively suggest that Hsp110s may not exhibit canonical nucleotide driven domain movements as seen in Hsp70s. At the same time, SAXS-based models of the Apo state deviate significantly from the SAXS based models of nucleotide bound states in specific areas of the protein. Furthermore, SAXS models of Apo state fitted best with PDB:2QXL (Liu & Hendrickson, 2007) where a small portion of SBDβ (residues 501-526 which is specifically found in Hsp110s but not in Hsp70 SBDs) is missing due to poor resolution. On the contrary, SAXS models of ATP or ADP states of Sse1 were best fitted with PDB:3C7N (Schuermann et al., 2008) which contains this region intact.

In order to understand this phenomenon clearly and also to realize the apo state at a higher resolution, we generated apo state of Sse1 using all-atomistic MD (Molecular Dynamics) simulation. The simulation revealed that the protein undergoes ~5 Å deviation from the starting structure within the first 100 ns (Figure S1A), we confirmed the stability of the protein for a long duration (~ 900 ns) (Figure S1B). MD stabilized final structure (Figure S1C) was found to be very similar to the nucleotide bound structure but with SBDβ stabilized and was folded within and not extended as seen in the ADP-BeFX-bound crystal structure of Sse1 (PDB ID:3C7N). We also found that SBDβ and SBDα regions were highly dynamic (Figure S1E, left and right panels). Finally comparing MD simulations with SAXS data suggest that nucleotide driven conformational changes occurs at the SBDβ (especially at the 500-540 residues) and SBDα resulting in unique sub-domain movements characteristic of Hsp110s (Figure S1E, left and right panels).

### Substrate peptide binding does not lead to closure of the α-helical lid of SBD of Sse1

Previous studies on multiple members of Hsp70s have shown that Hsp70 SBDs adopt significantly lid closed structure as a result of synergistic effect of substrate binding and ATP hydrolysis. In order to check if Sse1 adopts such Hsp70-like lid closed structure in the presence of substrate and ADP, we subjected Sse1 to SAXS measurements in the presence of ADP and LIC peptide (LICGFRVVLMYRF), a peptide substrate described for Sse1 previously (Figure 3A) (Goeckeler, Petruso et al., 2008). The Guinier analysis showed that the structural parameters remain relatively similar to that of apo and nucleotide bound states with a small increase in Rg to a value of 3.83 ± 0.16. However, comparison of P(r) profiles of Sse1 in substrate/ADP-bound state with ADP-only state showed that substrate binding increased the Dmax to 14.2 nm compared to 12.2 nm in case of ADP-Sse1 (Figure 3B). The protein continued to behave like a globular protein as shown by unchanged normalized Kratky plot with a peak at 1.73. Dummy residue models generated using GASBOR (Svergun, Petoukhov et al., 2001) showed SBD in open conformation as seen in apo and nucleotide-bound states (Figure 3C), except that the SBD lid undergoes slight structural changes which remain un-interpretable due to technical limitations. Thus we conclude that substrate binding does not lead to significant lid closure like canonical Hsp70s, rather it induces conformational changes that lead to increase in dimensions of Sse1, indicating unique mode of substrate binding by Sse1 or Hsp110s in general.

**Figure 3:**
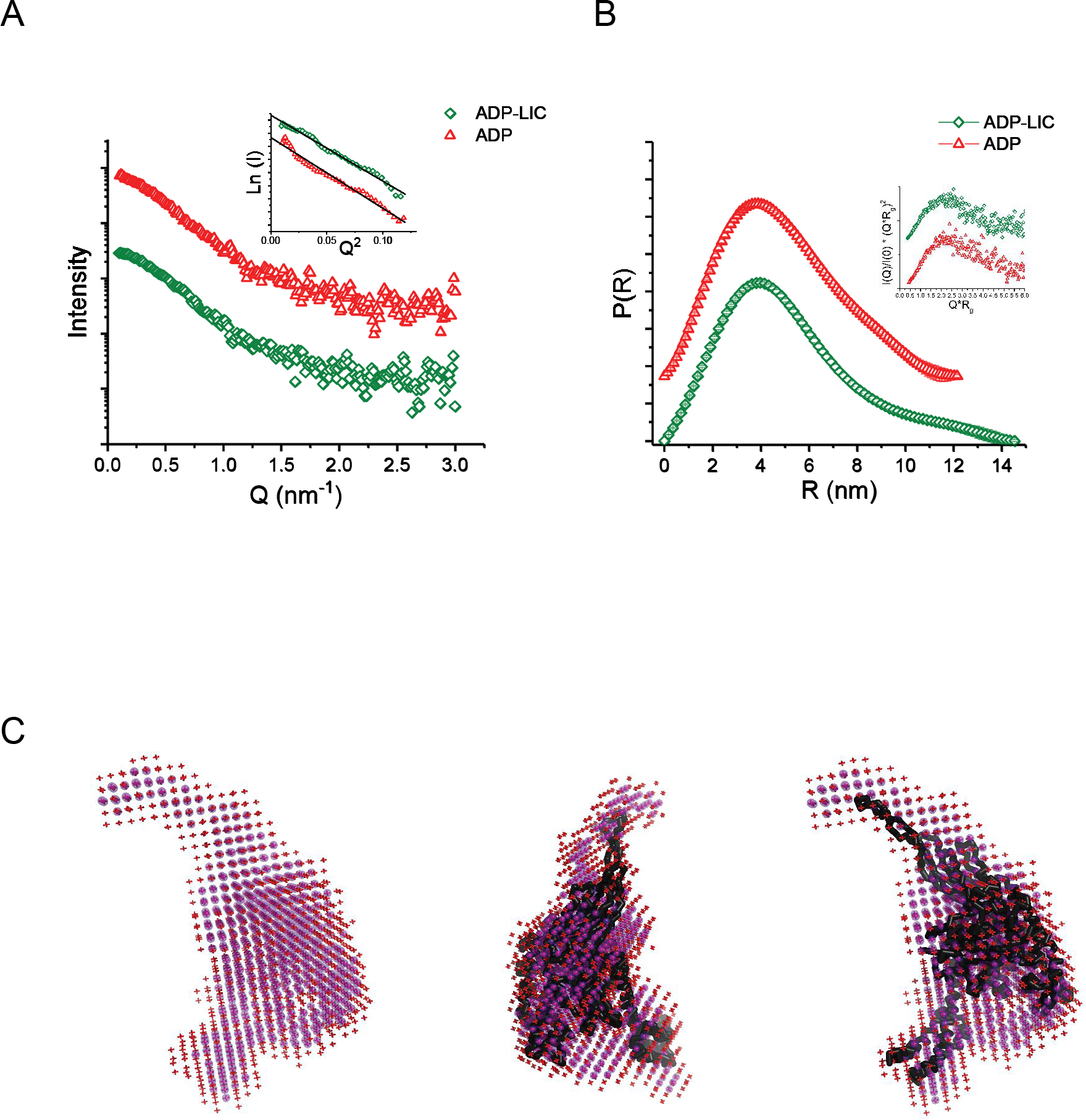
Peptide binding leads to conformational changes but not closing of the lid. **(A).** SAXS intensity profiles of Sse1 bound to the peptide substrate (LIC-Sse1) plotted as I(Q) (log scale) vs. Q (linear scale) with the SAXS profile of ADP-Sse1 added for comparison. The Guinier plots with the black solid lines showing the linear fits are shown in the inset. **(B)** The distance distribution curves (P(R) vs. R) for LIC-Sse1 with the normalized Kratky plots (I(Q)/I(0)*(Q*Rg)^2^ vs. Q*Rg) in the inset. Both plots have the corresponding profiles for ADP-Sse1 for comparison. **(C)** Starting from left, the damfilt (blue) and damaver (red) models for LIC-Sse1 followed by two orthogonal views of their overlay with the crystal structure of ATP bound structure (PDB ID: 2QXL).

### Hsp70 like domain conformation is recapitulated in chimeric constructs of Ssa1-Sse1

As the previous high-resolution structures of Sse1 (both in isolation and in complex with Hsp70 partners) have shown extensive inter-domain contacts between its NBD and SBD, we were curious to know the outcome of artificial fusion of the N-terminal domain of Hsp70 (NBD) and the C-terminal domain (SBD) of Hsp110 protein and vice versa. We asked the question whether the chimeric molecule would be stable as a two-domain protein. If it is stable, will the protein regain Hsp70-like domain movements or be static like Hsp110s? Is there a dominant effect of one of the domains in determining the nature of inter-domain allostery? There are reports in literature on chimeric yet functional Hsp70s (Makhoba, Burger et al., 2016, Shonhai, Boshoff et al., 2005, Strub, Rottgers et al., 2002, Suppini, Amor et al., 2004) and Hsp110s(Polier, Hartl et al., 2010), resulting from domain-shuffling between different Hsp70s or Hsp110s. These functional chimeric proteins proved that individual domain function as well as allosteric communication is maintained in a synthetic protein with similarly folded domains from foreign proteins. Considering the overall similar domain architecture of Hsp70s and Hsp110s, we proceeded to make chimeric constructs of these two different chaperone proteins. We made chimeric proteins by shuffling the domains of yeast Hsp110, Sse1 and its Hsp70 partner, Ssa1. Taking into consideration the importance of the inter-domain linker for solubility of NBDs of some Hsp70s (Blamowska, Sichting et al., 2010), we shuffled the NBDs with and without the respective linker sequences resulting in four different chimeric proteins. For simplicity, we have named them as Chimera 1(AAE), chimera 2 (AEE), chimera 3 (EEA) and chimera 4 (EAA) where the three letters denote the source of NBD, linker and SBD respectively from either Ssa1 (A) or Sse1 (E) (Figure 4A). All chimeras were expressed with cleavable N-terminal hexa-histidine tags in *E.coli* cells and we found that after induction at 30°C, chimera 1and 2 were mostly found in soluble fraction although chimera 3 and 4 were majorly found in the inclusion bodies (Figure S2A). When we expressed Chimera 3 and 4 in lower temperature (18°C overnight induction), solubility of these proteins were improved significantly and we were able to purify these proteins from the soluble fraction (Figure S2B). Expression of full length chimeric proteins were also checked in yeast cells by expressing them in *sse1*Δ cells under the endogenous promoter and immuno-blotting with polyclonal anti-Sse1 antibody raised against the full length Sse1 protein (Figure S2C). Indeed, all four chimeric proteins were expressed as full length proteins in yeast cells and could be detected by anti-Sse1 antibody. After we established that the chimeric proteins are stable, we were interested to study the domain arrangement of the chimeric proteins. To achieve that, we subjected chimera 2 (AEE) to SAXS measurements (Figure 4A). SAXS data analysis and modelling were done as described previously. The Guinier analyses revealed larger R_g_ of 4.25 ± 0.9 (Figure 4 A, inset) for Chimera 2 during its apo state and P(r) distribution profile showed larger D_max_ of 15.2 nm (Figure 4 B) compared to the wild type Sse1 protein. Also, the normalized Kratky plot of Chimera 2 had the peak shifted from the value of 1.73 indicating an increase in the flexibility of the protein (Figure 4 B, inset). Notably, SAXS model of chimera 2-apo could not be fitted to neither the Sse1-ATP structure (PDB code 2QXL(Liu & Hendrickson, 2007); NSD 3.7) nor to the Sse1-ADP*BeFX structure (PDB code 3C7N (Schuermann et al., 2008); NSD 4.8). On the contrary, it was best fitted with crystal structure of peptide-bound state of DnaK (PDB code: 2KHO; NSD 1.1)(Chang, Sun et al., 2008)(Figure 4C). This data suggests that the domain-undocked, lid-closed state of Hsp70s typically found in peptide-bound states of Hsp70s which probably has been lost in Hsp110s, can be artificially recapitulated in Hsp110, especially in its SBD. We tried to capture the structural features of other type of chimera where the NBD is obtained from Sse1 and SBD from Ssa1 (Chimera 3 and 4), at high protein concentrations required for SAXS measurements, both of these proteins were extremely aggregation prone and we were unable to obtain the data required to build a SAXS model.

**Figure 4:**
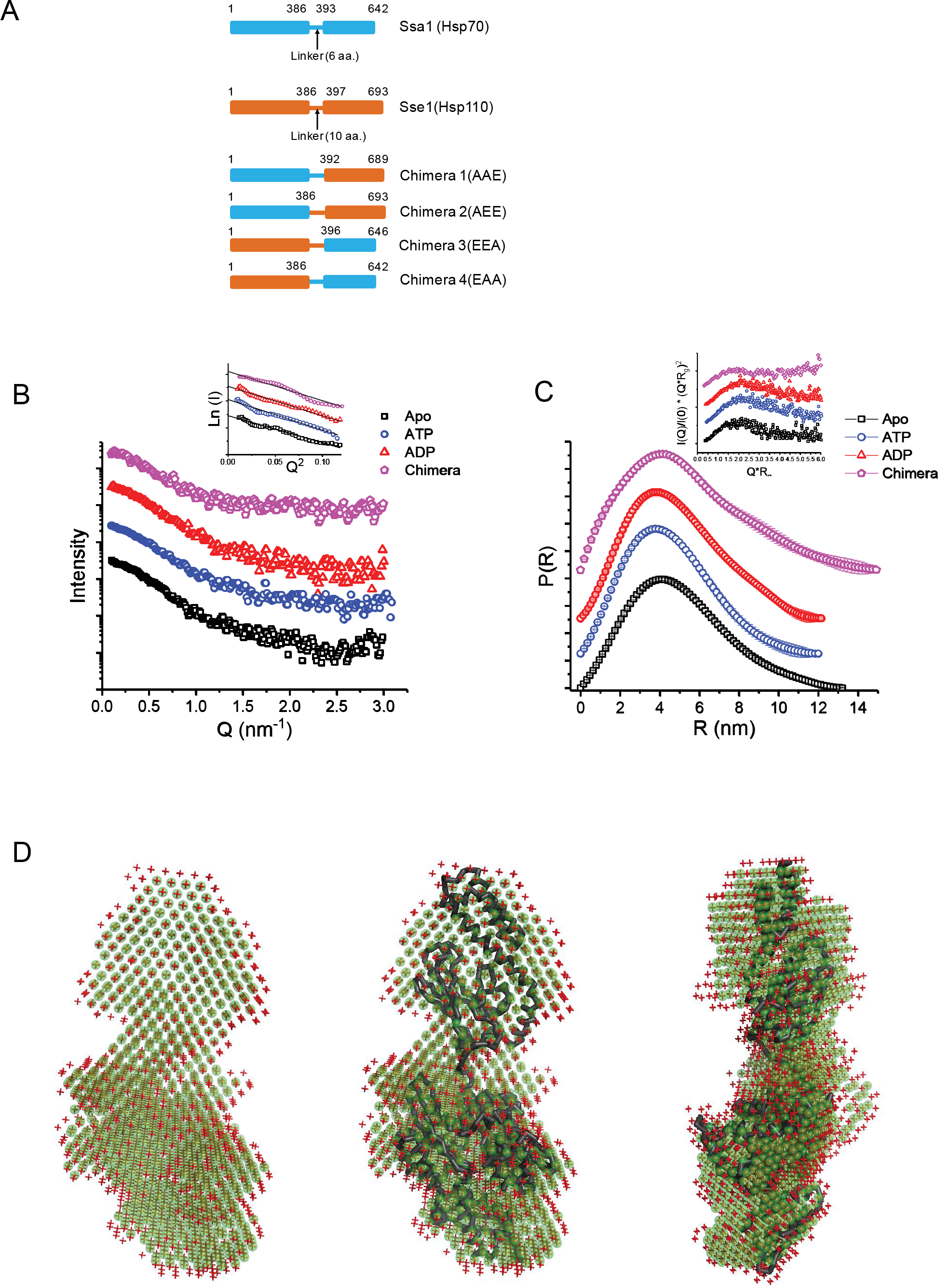
SAXS data analysis and modelling of the chimeric protein. **(A).** Schematic representation of chimeric constructs of Sse1 and Ssa1. The number denotes the amino acid numbers. **(B)** SAXS intensity profiles of the chimeric Sse1 (Chi-Sse1, Chimera 2) along with the SAXS profile of Apo-Sse1, ATP-Sse1 and ADP-Sse1 shown for comparison. The corresponding Guinier plot for the four datasets is shown in inset with the black solid lines showing the linear fits. **(C)** The distance distribution curves (P(R) vs. R) for Chi-Sse1, Apo-Sse1, ATP-Sse1 and ADP-Sse1 along with the normalized Kratky (I(Q)/I(0)*(Q*Rg)^2^ vs. Q*Rg) plots in the inset. **(D)** Starting from left, the damfilt (green) and damaver (red) models for Chi-Sse1 followed by two orthogonal views of their overlay with the crystal structure of peptide bound DnaK (PDB ID: 2KHO).

### Chimeric proteins of Ssa1-Sse1 are compromised of the nucleotide exchange factor function

As the chimeric proteins were stable and could be purified as full length two-domain proteins and additionally Chimera 2 artificially regained the Hsp70 like domain movements, it was interesting to check whether the chimeric proteins can replace the wild type Sse1 protein. Previous studies have shown that the double deletion of two paralogues of Hsp110 of yeast, *sse1* and *sse2* is synthetically lethal due to lack of efficient exchange factors for cytosolic Hsp70s. We checked whether the 70-110 chimera can rescue this synthetic lethality of yeast by chasing the Ura-plasmid containing wild type copy of Sse1 on 5-FOA (5-Fluoro-orotic acid) containing plates. When the chimeras were expressed from the endogenous promoter of Sse1, it could not rescue the synthetic lethal phenotype of double deletion of Sse1 and Sse2 indicating complete absence or insufficient NEF activity of 70-110 chimera (Figure 5A). To check whether over-expression of chimeras can rescue the phenotype, we further over-expressed the chimeras from high-copy number (2μ) plasmids. Importantly, over-expressing chimeras also could not rescue the synthetic lethal phenotype of double deletion of Sse1 and Sse2 although over-expressing Fes1, a cytosolic NEF, could rescue the lethal phenotype as described earlier (Figure 5B) (Raviol, Sadlish et al., 2006). This result led to the conclusion that although the chimeric proteins are expressed as stable two-domain full length proteins, they are not able to perform the NEF function possibly due to regaining of Hsp70 like domain movements which may be detrimental for the NEF activity of the Hsp110 proteins.

**Figure 5:**
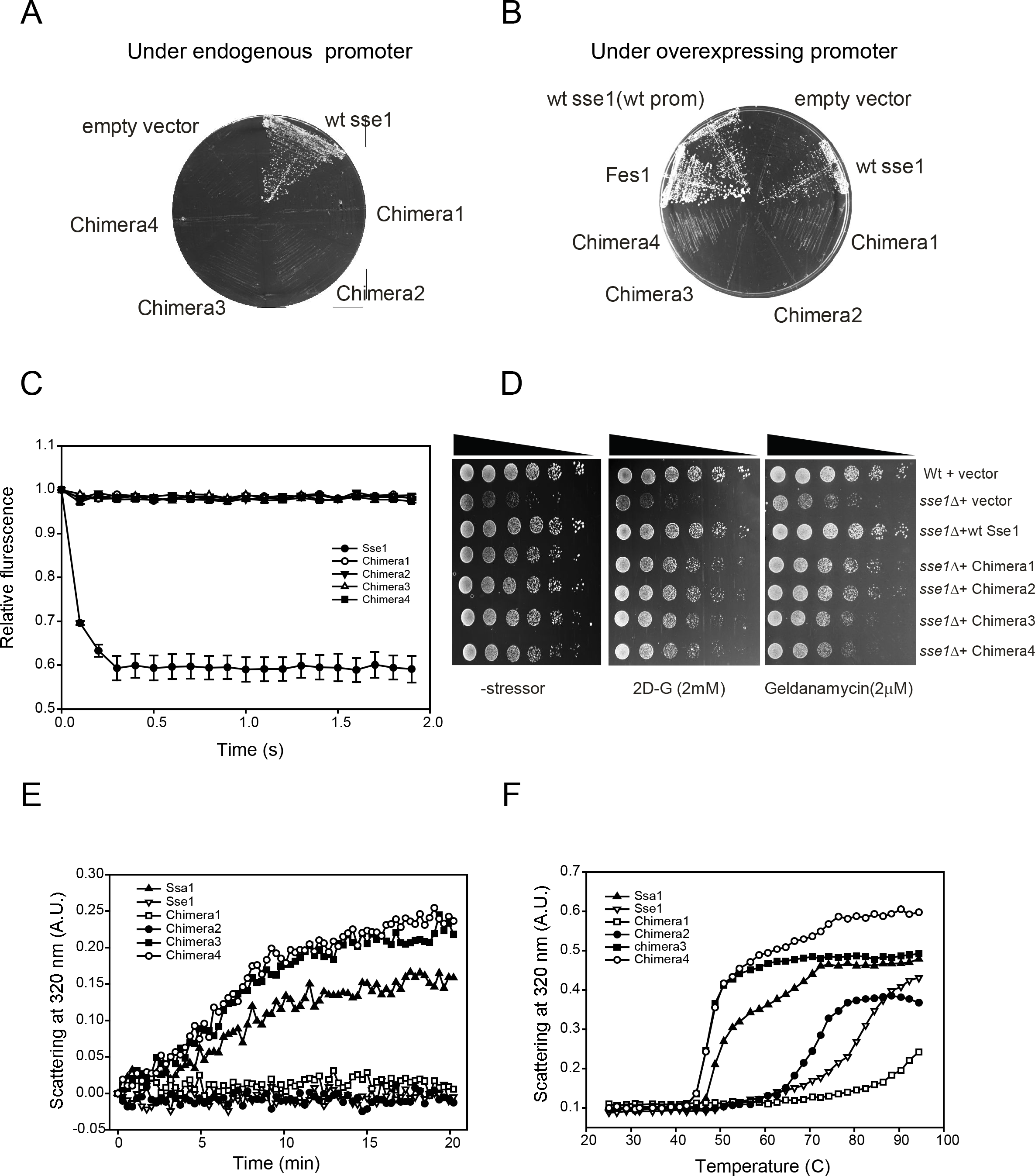
Ssa1-Sse1 chimeric constructs are compromised of the nucleotide exchange factor function. The endogenous **(A)** and over expression **(B)** of wild-type Sse1 and chimera in sse1Δ-sse2Δ strain (sse1::His3 sse2::KanMX4 pCM189-SSE1-URA3, generous gift from Prof. F.U. Hartl’s group from MPI, Biochemistry) shows that chimera are not capable of rescuing the lethal phenotype of sse1Δ-sse2Δ whereas the overexpression of another cytosolic nucleotide exchange factor, Fes1 can rescue sse1Δ-sse2Δ phenotype. Panel **C** shows the nucleotide exchange assay for the wild-type and chimera by the measuring the decrease in the fluorescence of Mant-ATP upon release due to exchange, which reveals that wild-type Sse1 efficiently exchange the nucleotide whereas all four chimera lack the nucleotide exchange factor activity. (**D)**. Growth phenotype of sse1Δ cells are rescued by all four chimera like wild-type *sse1* under physiological condition although when the cells are subjected to proteotoxic stressors (geldanamycin and 2- Deoxy glucose), by chimera 1 and 2 exhibit appreciable growth rescue compared to wild-type sse1 whereas chimera 3 and 4 exhibit severe growth defects. (**E)**. Aggregation propensity of chimeric proteins were measured by subjecting the wild-type and chimera for thermal denaturation at 42°C and following the light scattering at 320 nm. Chimera 3 and 4 along with Ssa1 exhibited increased light scattering with time (20 min) indicating aggregation whereas chimera 1, 2 and wild-type Sse1 did not exhibit any increase in light scattering at 42°C indicating no thermal aggregation at this temperature. (**F)**. Chimera and wild-type Sse1 and Ssa1 proteins were subjected to thermal melting from 25° to 95°C by following the light scattering at 320 nm. Ssa1 and chimera 3 and 4 exhibit appreciable light scattering at ~45°C indicating aggregation whereas Sse1 and chimera 1 and 2 are much more thermally stable and shows aggregation beyond 65°C only.

To find any other alterations in the domain-functions in the chimeric proteins from the parent protein from which they are obtained, domain functions were determined by measuring the ATP-binding (FIGS3) and substrate binding activity of NBD and SBDs respectively (Table S2). All chimera exhibited ATP binding and comparable affinity to the LIC peptide (Goeckeler et al., 2008) as measured by SPR measurements (Table S2). As the nucleotide binding and substrate binding activities of chimera were comparable to its parent proteins, it was interesting to examine whether the chimeric proteins harbor any residual exchange factor function, as it was expected from the in vivo experiments that either the chimeras are completely deficient as NEF or are only partially active and are insufficient to rescue the synthetic lethal phenotype of double deletion of Sse1 and Sse2. All four recombinant chimera, purified from *E.coli*, were checked for nucleotide exchange activity by following the change in fluorescence signal of MANT-ATP upon release from Hsp70s after being exchanged with non-fluorescent nucleotides. Wild-type Sse1 showed prominent exchange activity while none of the chimera showed any exchange activity (Figure 5C). This result points towards the fact that important inter-domain communication or interactions are critical for NEF activity of Sse1 (Hsp110) proteins.

### Ssa1-Sse1 chimeric proteins are effective as chaperones during proteotoxic stress

Apart from its’ co-chaperone function as NEF, Hsp110s have been implicated in many other cellular functions, especially restoration of protein homeostasis under proteotoxic threats in various eukaryotic models, although Hsp110’s role in such processes as chaperone is still not clear (Kuo et al., 2013, Zhang et al., 2010). Thus, it was interesting to check whether the chaperone functions of Sse1 can be performed by the chimeric proteins. For that purpose, all chimeric proteins were cloned under endogenous promoter of Sse1 retaining the 3’UTR of Sse1 in centromeric plasmids and the plasmids were transformed in *sse1*Δ cells. Wild-type Sse1 was cloned similarly and transformed in *sse1*Δ cells to serve as positive control and the empty plasmid was transformed as the negative control. All the resulting *sse1*Δ yeast strains harboring plasmids containing the chimera were grown under permissive (30°C) as well as in media containing small molecules like Geldnamycin (an inhibitor of Hsp90) or 2-D-G (an inhibitor of glycolysis) that perturb the protein homeostasis and have been shown to be toxic for *sse1*Δ cells (Liu et al., 1999). When the growth of *sse1*Δ cells were compared with the wild type cells by drop-dilution assay, prominent growth defect of *sse1*Δ cells were evident both in presence and absence of toxic chemicals (Figure 5D). These growth phenotypes were completely rescued in the cells expressing wild-type Sse1 from plasmids. Interestingly, chimera 1 (AAE) and chimera 2 (AEE) showed growth rescue like the wild type Sse1 protein in all conditions.

In contrast, chimera 3 (EEA) and chimera 4 (EAA) could rescue the growth phenotype albeit much less significantly (Figure 5D). To find out the reason of differential rescue potential of *sse1*Δ phenotypes by two groups of chimera, we checked its’ expression levels. Indeed, the protein levels of chimera 3 and 4 were less compared to chimera 1 and 2 and this may be a possible explanation of less efficient rescue of *sse1*Δ phenotypes by chimera 3 and 4. We reasoned that the less abundance of chimera 3 and 4 proteins could be due to compromised stability of these proteins. In conclusion, we demonstrate that growth phenotype of *sse1*Δ cells under proteotoxic stresses are completely rescued by the chimera containing the Ssa1 NBD and Sse1 SBD while the growth phenotype was rescued only partially by chimera 3 and 4 harboring Sse1 NBD and Ssa1-SBD.

### The SBD of Sse1 exerts chaperoning activity on self NBD and structurally similar foreign NBDs

To corroborate our *in vivo* observation that chimera 3 and 4 were less efficient compared to chimera 1 and 2 in rescuing the growth defects of *sse1*Δ cells under proteotoxic stresses, plausibly due to low stability, we performed *in vitro* stability experiments. We measured the aggregation propensity of these chimeric proteins along with wild-type Sse1 and Ssa1. We incubated all chimeric proteins and the wild type Ssa1 and Sse1 proteins at 42°C and monitored the increase in light scattering at 320 nm as an indicator of aggregate formation during the thermal denaturation of proteins at 42°C (Figure 5E. As shown in Figure 5E, the scattering for chimera 1, chimera 2 and wild type Sse1 remained in the baseline level indicating absence of aggregation upon subjecting the proteins to high temperature at 42°C. In contrast, chimera 3 and 4 showed significant increase in scattering within 5 minutes of incubation at 42°C indicating aggregation. Interestingly, wild type Ssa1 also demonstrated significant scattering albeit appreciably less compared to scattering exhibited by chimera 3 and 4. This experiment hinted that chimera 3 and 4 are unstable and aggregate at 42°C due to thermal denaturation whereas chimera 1 and 2 are more stable at this temperature. To get an idea about the difference in thermal stability between these two types of chimeric proteins, we subjected all protein to thermal melting where we heated the proteins gradually from 25°C to 95°C while monitoring the scattering at 320 nm wavelength. As shown in Figure 5F, chimera 1 and 2 are significantly thermally stable like wild type Sse1 protein and starts aggregating only beyond 65°C. In contrast, chimera 3 and 4 are much less stable and starts aggregating at 45°C only. Interestingly, we found wild type Ssa1 to be much less stable compared to Sse1 during thermal denaturation. In summary, we observed that chimeric proteins with Ssa1 NBD and Sse1 SBD are significantly more stable like wild type Sse1 than the chimera containing Sse1 NBD and Ssa1 SBD. Our data demonstrate that Sse1 SBD has a strong chaperoning capacity which helps to stabilize not only its own NBD as exhibited by great thermal stability of wild type Sse1 protein but also NBD of similar architecture of foreign proteins like Ssa1 (as shown by substantial thermal stability of chimera 1 and 2).

### The NBD of Sse1 is important for substrate peptide binding

Although Hsp110 chaperones are considered as structural homologues of Hsp70 chaperones, few alterations in structure are easily discernible in Hsp110 (Sse1) especially in its SBD. The implication of such alteration in structure of SBD on substrate binding or chaperoning capacity of Hsp110, remain underexplored. In case of Hsp70s, isolated SBDs retain the substrate binding properties almost at the near-native level and the substrate binding pocket is harboured in the β-sheet domain. In case of Hsp110s, the position of substrate binding pocket is still enigmatic. In light of our current finding that the domains are relatively rigid with respect to each other, with minor conformational changes in presence of nucleotides, it was interesting to investigate the exact structural features required for peptide binding by Hsp110. When we analyzed the binding affinity to already described LIC peptide (Goeckeler et al., 2008), we found that the affinity of chimera 4 is equivalent to wild type Sse1 (Table S2) and that of chimera 2 is ~10 fold lesser. This finding was interesting because chimera 4 contains the SBD of Ssa1 and supposed to bind the LIC peptide with less affinity if the substrate binding pattern follows canonical binding mode as in case of Hsp70s (Rudiger, Germeroth et al., 1997). In contrast, chimera 2 contains the Sse1 SBD and is expected to bind like wild type Sse1 assuming similar peptide binding behavior of Hsp110 SBDs. To check for overall binding specificities of chimera in comparison to wild type Sse1, we probed the binding of wild type Sse1, chimera 2 and chimera 4 to a tiling array of overlapping peptides of Firefly luciferase (F-luc) protein. As shown in Figure 6A, wild type Sse1 shows prominent binding to few peptides in the pep-spot membrane as shown in the Table 1. Importantly, the peptides at spot numbers 87 and 88 demonstrating strongest binding to Sse1, shares the overlapping stretches of amino acids with the LIC peptide. In our F-luc pepspot array, LIC peptide constitute the 86^th^ spot but surprisingly we did not observe any noticeable binding to this spot although we observed significant affinity to this peptide by SPR as described before. Interestingly, chimera 2 (AEE) showed similar binding pattern to the F-luc pepspot membrane as found with wild type Sse1 although much less significantly with most of the spots (Figure 6B). This result again pointed out that chimera 2 in spite of harboring the SBD of Sse1, exhibits lesser binding affinity to same peptides indicating important role of NBD in efficient substrate binding. Intriguingly, chimera 4 (EAA) showed similar binding pattern and was bound to the spots with similar or even higher affinity on the membrane, especially the spots at positions 87-91 were most prominently bound to chimera 4 (Figure 6C). As chimera 4 harbors the SBD from Ssa1, we performed the pepspot binding assay with wild type Ssa1 protein which exhibited completely different binding pattern (Figure 6D) nullifying the possibility of enhanced binding by Ssa1-SBD in Chimera 4 to peptide spots with high affinity. This result explained the similar affinity of chimera 4 and Sse1 to LIC peptide by SPR measurements as described earlier. This data conclusively pointed out the importance of Sse1 NBD in binding the peptide substrates. It is tempting to speculate that during the course of evolution, the substrate binding site of Hsp110s had shifted from SBD to NBD-SBD interface and SBD has gained additional chaperoning or stabilizing function.

**Figure 6:**
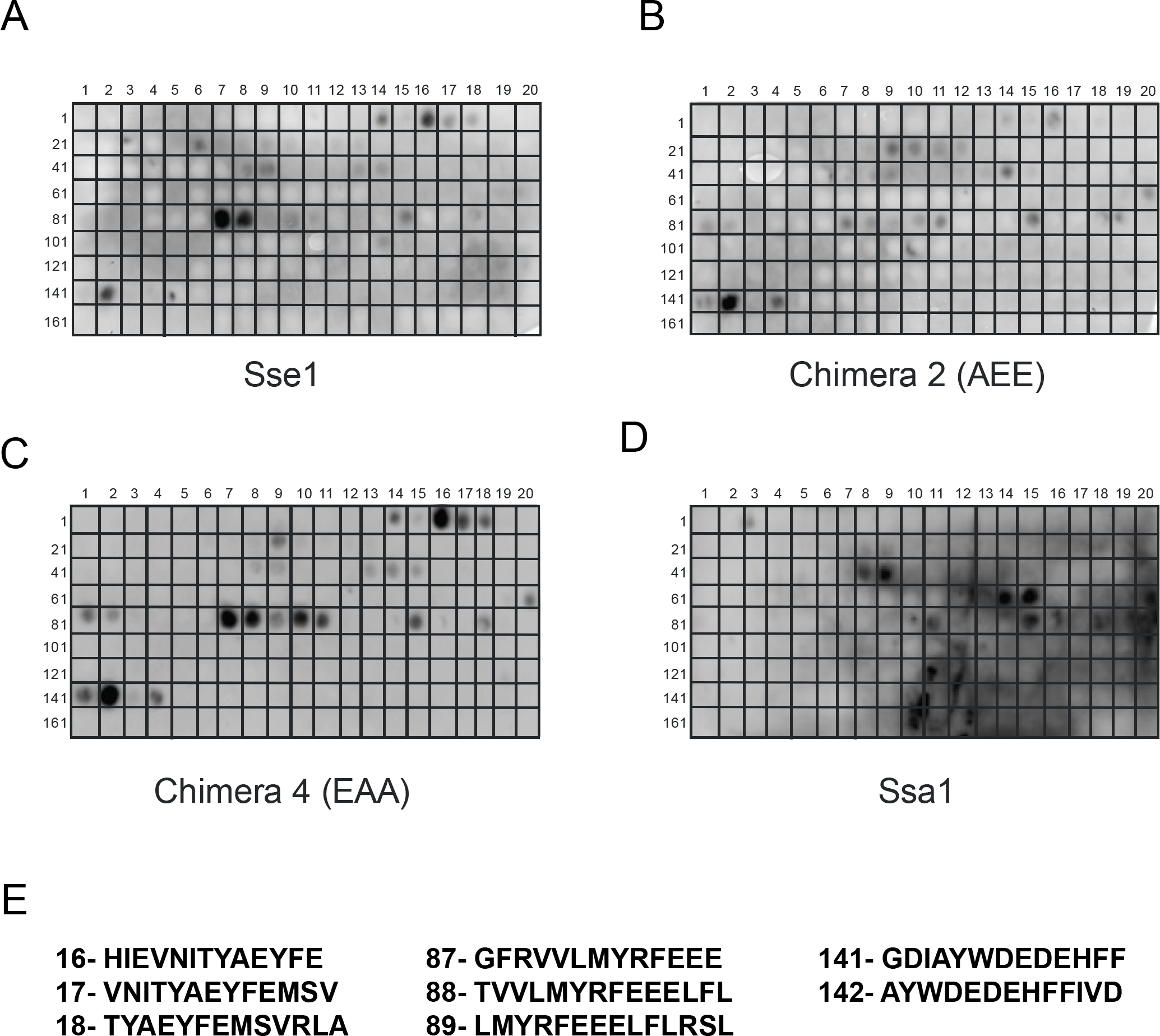
The Nucleotide Binding Domain of Sse1 is important for substrate peptide binding. The peptide binding signature of wild-type and chimera were determined using the Firefly Luciferase Peptide array containing overlapping 13mer peptides from F-luc immobilized on the cellulose membrane (JPT peptides, GmbH). **(A-D)** The peptide binding signature of wild-type Sse1, chimera 2, chimera 4 and wild-type Ssea1, respectively. Panel A shows the wild-type Sse1 binding to peptide array in the pep-spot membrane. The peptide spots 87 and 88 show strongest binding to Sse1 and these peptides share the overlapping sequence with the LIC peptide. B. The peptide scan for chimera 2 (AEE) with the F-Luc peptide array shows lower affinity to the peptide spots bound to wild type Sse1 whereas chimera 4 (EEA) **(C)** shows stronger affinity towards the Sse1 interacting peptides. Both chimera 2 and 4 show distinct peptide binding patterns. On the contrary, Ssa1 has completely distinct binding signature **(D)**. Panel **E** shows the peptide sequences of the significantly-bound peptide spots.

**Table 1.**
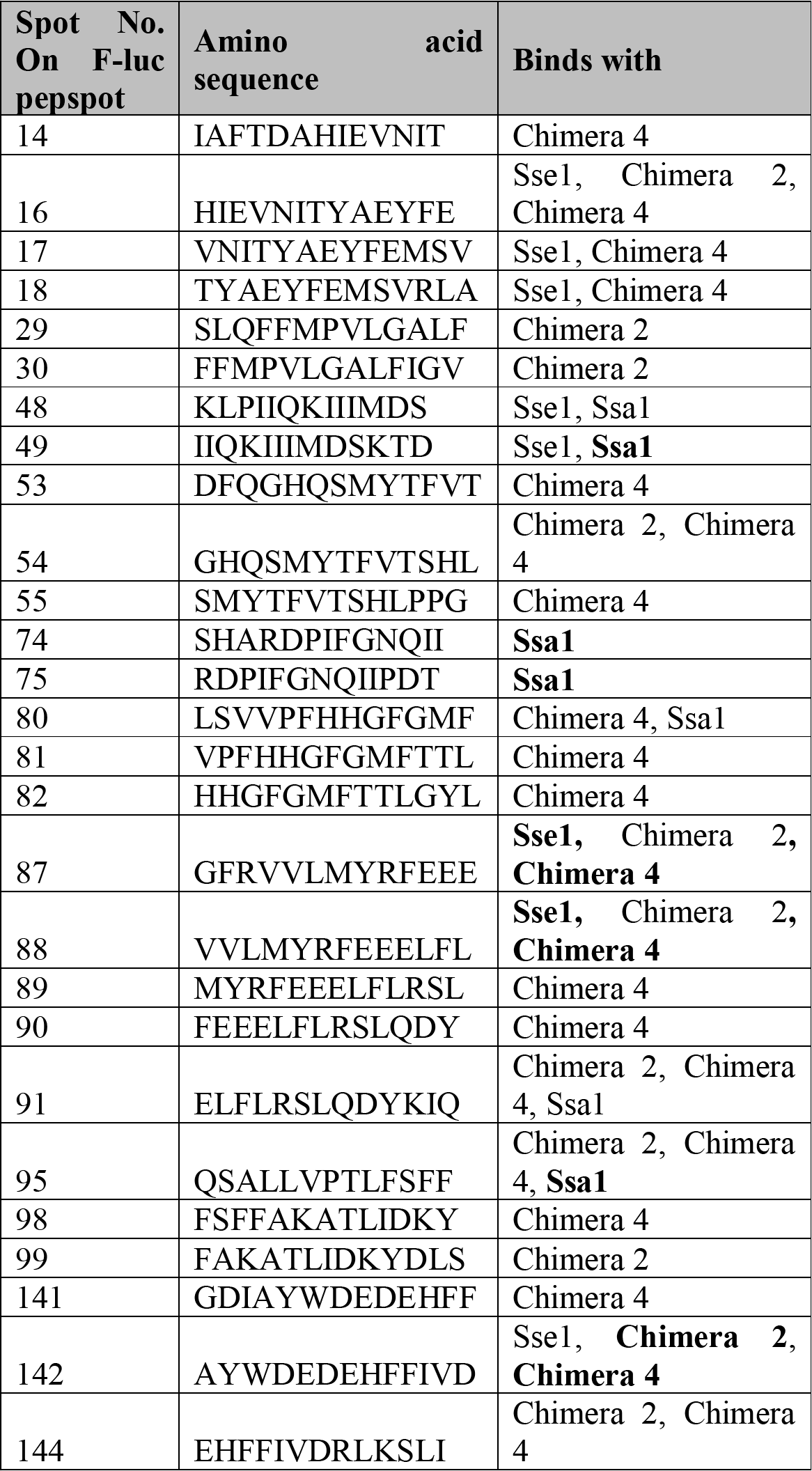
The sequences of all peptides on the pep-spot array to which wild-type Sse1, Ssa1 and the chimeric proteins exhibited appreciable binding, have been tabulated. The proteins written in ‘bold’ fonts showed significant binding to the respective peptides.

## Discussion

In the current study, we present unconventional domain movements and domain functions of Hsp110 proteins taking yeast Hsp110, Sse1 as a model. Although the high resolution structure of Sse1 was solved about a decade ago, the dynamic nature of the domains that may be crucial for cellular functions of these group of molecular chaperones, remained enigmatic. By employing two different state of the art techniques, single molecule FRET spectroscopy and Small-angle X-ray scattering (SAXS), we have captured, unique conformational changes of the Hsp110 protein in solution. Furthermore, we have done MD simulation to observe finer changes which were difficult to capture by SAXS. Indeed, using a combination of various techniques, we conclusively demonstrate that despite significant similarity in domain organization and individual domain structures, Hsp110s are unique and deviated significantly from its structural homologue, Hsp70s. The most prominent and well described conformational changes in Hsp70s are opening-closing of α-helical lid of SBD and the docking-undocking of two domains, NBD and SBD during the ATP-state and ADP/substrate-bound state, respectively. So far, only ATP or ATP-analogue bound states of Hsp110 were captured in high resolution leaving behind a caveat in understanding of the structural features of the protein in its other functional states. Here, using combination of two different methodologies, sm-FRET and SAXS, we show that indeed, Hsp70-like lid movements or inter-domain communication are lacking in Hsp110. We show that α-helical lid remains wide open in ATP, ADP or even substrate bound states although there are additional subtle changes in the β-sheet base of SBD and the α-helical lid as captured by SAXS models and nicely recapitulated by MD simulation. This data indicates that the mechanism of substrate binding and the position of substrate binding pocket may be completely different for Hsp110s compared to canonical Hsp70s. This is further supported by our substrate binding experiments with SPR measurements with a single substrate peptide or with an array of peptides. In both cases, we demonstrate that the Hsp70-110 chimeric proteins harboring 110-NBD recapitulates the binding specificity and affinity of the wild type Hsp110 rather than the ones harboring its SBD. This is completely counterintuitive considering the substrate binding mechanism of canonical Hsp70 chaperones where the isolated SBDs are nearly identical in substrate binding capacity to full length proteins. On the other hand, we show that SBD has an inherent stabilizing capacity to self as well as structurally similar NBDs. In summary, our data reveal a completely new insight into the evolution of Hsp110 domain communication and functions.

## Methods

### Strains and Plasmids

Wild-type *sse1*, *ssa1* and chimeras were cloned in the pETDuet1, pRS313 and pJV340 vector for propagation in *E. coli* and *S. cerevisiae*. The Chimeras were constructed by fusing the domains of sse1 and ssa1 by overlap PCR. These Constructs and wildtype sse1, ssa1 were inserted into the pETDuet1 using the restriction sites BamH1/Sal1 or Sac1 for sse1 and ssa1 respectively. The yeast centromeric plasmids (pRS313 and pRS316) were inserted with the Sse1 Promoter and 3’ UTR using the restrictions site Sac1/BamH1 and Xho1/Kpn1 respectively. Further, we introduced wild type sse1, ssa1 and chimeras in between Promoter and 3’ UTR using the restriction sites BamH1/Xho1. And also the wild-type Sse1 and chimeras were cloned into the Yeast 2μ plasmid (pJV340) using the restriction sites BamH1/Xho1. All the constructs were confirmed by sequencing. Yeast cells were cultured in the selective synthetic medium at 30^o^C for growth and maintenance of plasmids. For drop dilution assay, yeast cells were serially diluted from 1 O.D and equal volume were spotted on selective synthetic medium with or without proteotoxic stress.

### Purification of Proteins

For purification of the His_6_ tagged wild-type Sse1, Ssa1 and chimeras, we expressed them in the *E.coli* BL21 cells with IPTG induction (final 0.5mM) at 18^°^C, overnight. The overnight grown cells were harvested and the cells were lysed in the Buffer A with 1mM (final) PMSF (Buffer A:25mM HEPES (pH 7.4), 250mM KCl, 1mM β–Mercaptoethanol, 10mM Imidazole). The lysate was incubated with Ni^2+^ NTA resin (Sigma). The bound proteins were eluted with Buffer A+ 250mM Imidazole. The Eluted proteins were buffer exchanged with storage buffer (25mM HEPES (pH 7.4), 150mM KCl, 5mM MgCl_2_, 1mM β–Mercaptoethanol, 10% Glycerol) and aliquots of protein were stored in -80^O^C.

### ATP binding and Nucleotide exchange assay

The ATP binding activity of the proteins were measured by incubating 100nm of Mant-ATP (Anaspec) with 1μM of protein (wild-type Sse1/Ssa1/Chimeras) in buffer containing 25mM HEPES (pH 7.4), 150mM KCl and 5mM MgCl_2_. The fluorescence emission spectra from 380 to 600nm were recorded by spectrofluorimeter (Horiba Fluorolog-3) following exciting the mant-ATP at 355nm. Controls where the fluorescence of MANT-ATP alone and buffer alone were measured to compare the effective binding of ATP to protein of interest. The stop flow measurement of MANT ATP was measured in Horiba Fluoromax-3. The nucleotide (ATP) exchange rate of 1μM Ssa1 were measured by exchange of 100nm MANT ATP with 200mM ATP in the presence of 1μM sse1 and all four chimeras.

### Surface Plasmon Resonance

The dissociation rate constants for wild type Sse1 and chimera’s interaction with LIC peptide was measured by SPR measurements by BIAcore3000. LIC peptide (100mM) was immobilized on CM5 amine coupling chip according to manufacturer’s protocol using immobilization buffer (10mM sodium acetate, pH 4.5). Before the run, the peptide-immobilized lane and the reference lane (blank) were equilibrated with BIAcore assay buffer (25mM HEPES, 150mM KCl, 5mM MgCl2, pH 7.4). Wild type Sse1 and chimeras were passed over the peptide-immobilized chip at five different concentrations (10nM, 50nM, 100nM, 500nM, 1000nM) in BIAcore assay buffer and response was recorded. SPR data were exported to BIAevaluation tool and the response curves were fitted to Langmuir’s equation for calculating the equilibrium dissociation constant (KD) for wild type Sse1 and chimeras.

### Small Angle X Ray Scattering (SAXS) data collection and analysis

SAXS measurements were done on SAXSpace system (Anton Paar GmbH, Austria) with sealed tube (line collimation) X-ray generator (λ = 1.5418) operated at 40 kV and 50mA mounted with 1D CMOS MYTHEN (Dectris, Switzerland) detector. The scattering intensity I(q) was measured for scattering angles (Q = 4πsinθ/λ) from 0.01 to 3 nm^-1^. The samples were placed in a quartz capillary on a thermostatic capillary holder. The data was collected at 25°C and the integrated exposure time for one acquisition was 45 minutes containing 3 frames each of 15 minutes. Approximately 50μL of each sample was exposed to X-rays. Unless specified otherwise, the purified concentrated proteins were dialyzed against 20mM HEPES, 150mM KCl, 5mM MgCl_2_ and 2mM DTT at a final pH of 7.2. Immediately prior to SAXS experiments, samples were centrifuged in a tabletop centrifuge, and the protein concentrations were determined using a Nanodrop spectrophotometer (Thermo Scientific). SAXStreat software was used for data reduction and calibration of primary beam position. Data processing, including buffer subtraction, scaling and de-smearing was done using SAXSquant software. Estimation of radius of gyration (Rg), pair-wise distribution function (P(r)) and D_max_ from the SAXS profile of the protein samples was done using PRIMUSQT available in ATSAS suite (Franke, Petoukhov et al., 2017). To obtain solution structures, modelling of low resolution structures was carried out using DAMMIF(Franke & Svergun, 2009). Ten independent Ab-initio models were generated and averaged and filtered using DAMAVER suite of programs (Volkov & Svergun, 2003). All high resolution structures were overlaid with SAXS-derived model using SUPCOMP20 (Kozin & Svergun, 2001). Molecular graphical representations were made using PyMOL (The PyMOL Molecular Graphics System, Version 1.8, Schrödinger, LLC) ad UCSF Chimera 1.12 (Pettersen, Goddard et al., 2004). The IUCr guidelines published by Jacques *et al.* were followed for reporting SAXS data parameters (Jacques, Guss et al., 2012).

### Establishing the structure of the apo-form of Sse1 protein by MD simulation

To elucidate the high-resolution apo-form of the protein, molecular dynamic simulation was performed on the nucleotide-removed form of Hsp110 (PDB ID: 3C7N) using GROMACS 4.6.1 (Van Der Spoel, Lindahl et al., 2005). All atomistic simulations were carried out using the CHARMM36 all-atom force field (November release) (Brooks, Brooks et al., 2009, Denning, Priyakumar et al., 2011, Huang & MacKerell, 2013) using periodic boundary condition and TIP3P water model, developed by Jorgensen *et al.*, was used to solvate the protein and ions (Jorgensen & Jenson, 1998). The starting model was solvated in a periodic box with 97916 TIP3 water molecules. 26 Na^+^ ions were added to the solvent to neutralize electrical net charge of the protein. The system was then minimized for 50,000 steps using a steepest decent algorithm with a emtol of 200 KJ/mol after a minimization with emtol of 100 KJ/mol. This was followed by an equilibration run of 100ps in NVT ensemble with restrains on the protein atoms. The NPT ensemble was used for production simulation. Systems were simulated at 310K, maintained by a Berendsen thermostat with a time constant of 1 ps with the protein and non-protein molecules coupled separately. Pressure coupling was done employing a Parrinello-Rahman barostat using a 1 bar reference pressure and a time constant of 2.0 ps with compressibility of 4.5e-5 bar using isotropic scaling scheme. Electrostatic interactions were calculated using the Particle Mesh Ewald (PME) summation. The production run was performed for 1 microsecond.

We then used MD stabilized final structure as an input for an iterative SAXS-guided refinement using SREFLEX (Panjkovich & Svergun, 2016). The final model was validated using CRYSOL (Svergun, Barberato et al., 1995), which gave a chi-square fitting score of 1.094 to the apo- SAXS data (Figure S1D).

## Acknowledgements

VK acknowledges CSIR-Senior Research Fellowship. KM acknowledges SERB YSS grant (YSS/2015/000532) for funding, Shiv Nadar University for infrastructural support. We acknowledge Kausik Chakraborty and Asmita Ghosh for their critical comments.

## Supplementary Information

**Atypical domain communication and domain functions of a Hsp110 chaperone**

Vignesh Kumar^#^ ^1,2^, Joshua Jebakumar Peter^#^ ^1^, Amin Sagar^3^, Arjun Ray^1, 2^, Ashish ^3^* and Koyeli Mapa^2,4^*.

1. Proteomics and structural biology unit, CSIR-Institute of Genomics and Integrative Biology, Mathura Road, New Delhi 110025, India.

2. Academy of Scientific and Innovative research (AcSir), CSIR-CRRI, Mathura Road, New Delhi 110025, India.

3. CSIR-Institute of Microbial Technology, Sector 39 A, Chandigarh, 160036.

4. Department of Life Sciences, School of Natural Sciences, Shiv Nadar University, NH91, Greater Noida, Gautam Buddha Nagar, UP 201314, India.

*Address for correspondence,

#: contributed equally for this work.

## Supplementary table legends

**Supplementary Table 1.**
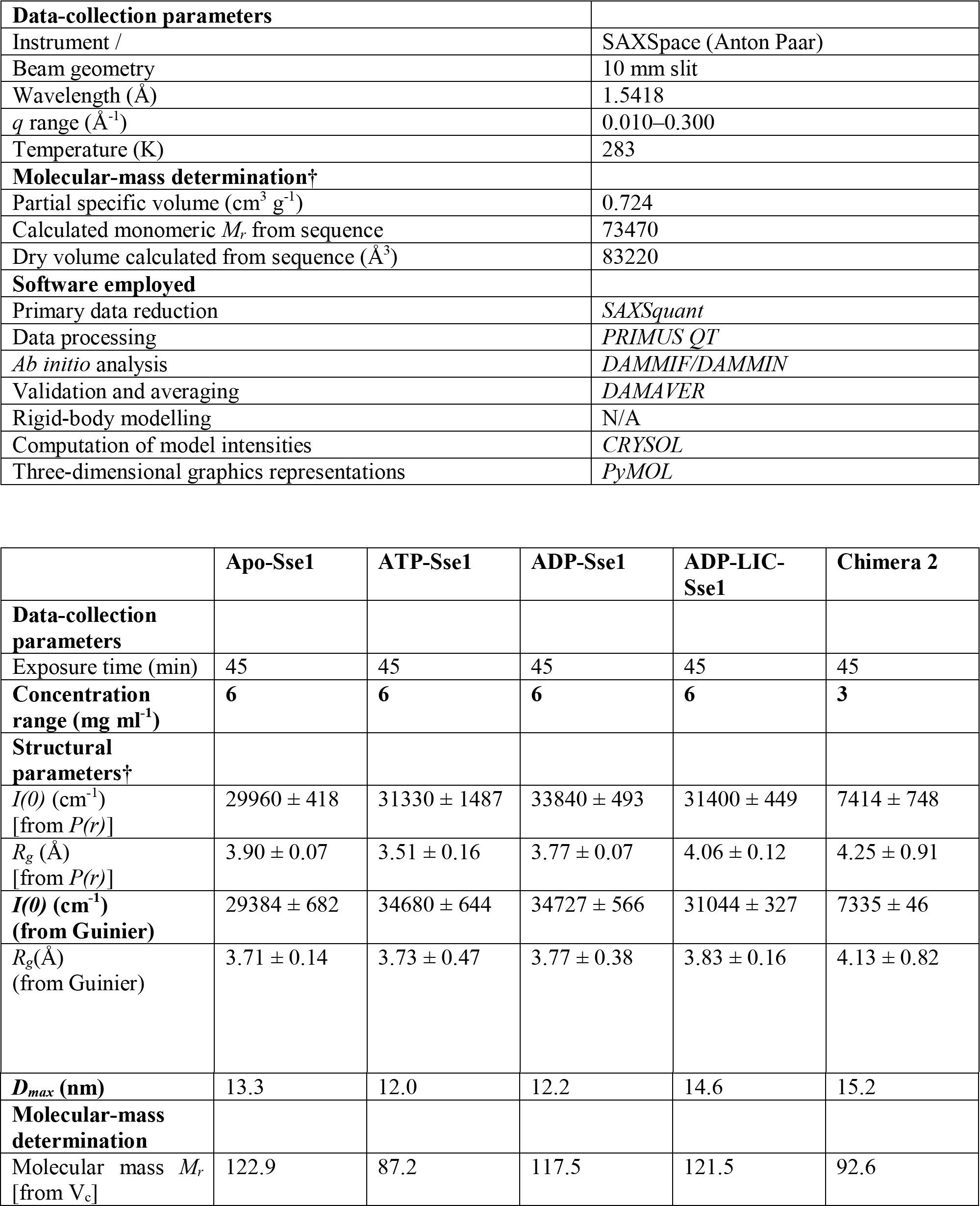
SAXS data-collection and scattering-derived parameters.

**Supplementary Table 2.**
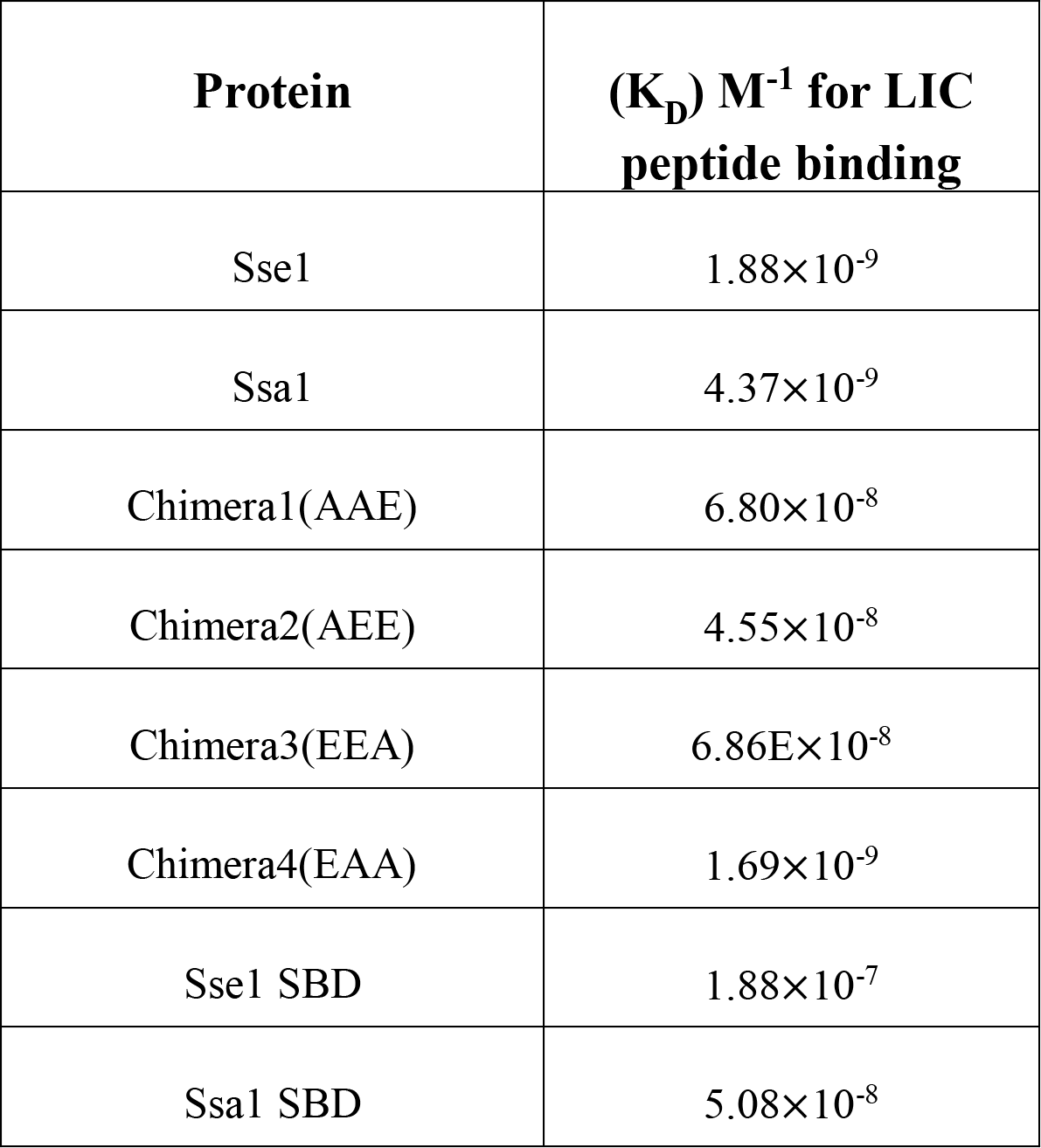
SAXS data-collection and scattering-derived parameters. The equilibrium dissociation constant (K_D_) for interaction of LIC peptide with wild-type Sse1 and Ssa1 were calculated as described in the methods section and have been tabulated in Table S2.

## Supplementary Figure legends

**Figure S1.**
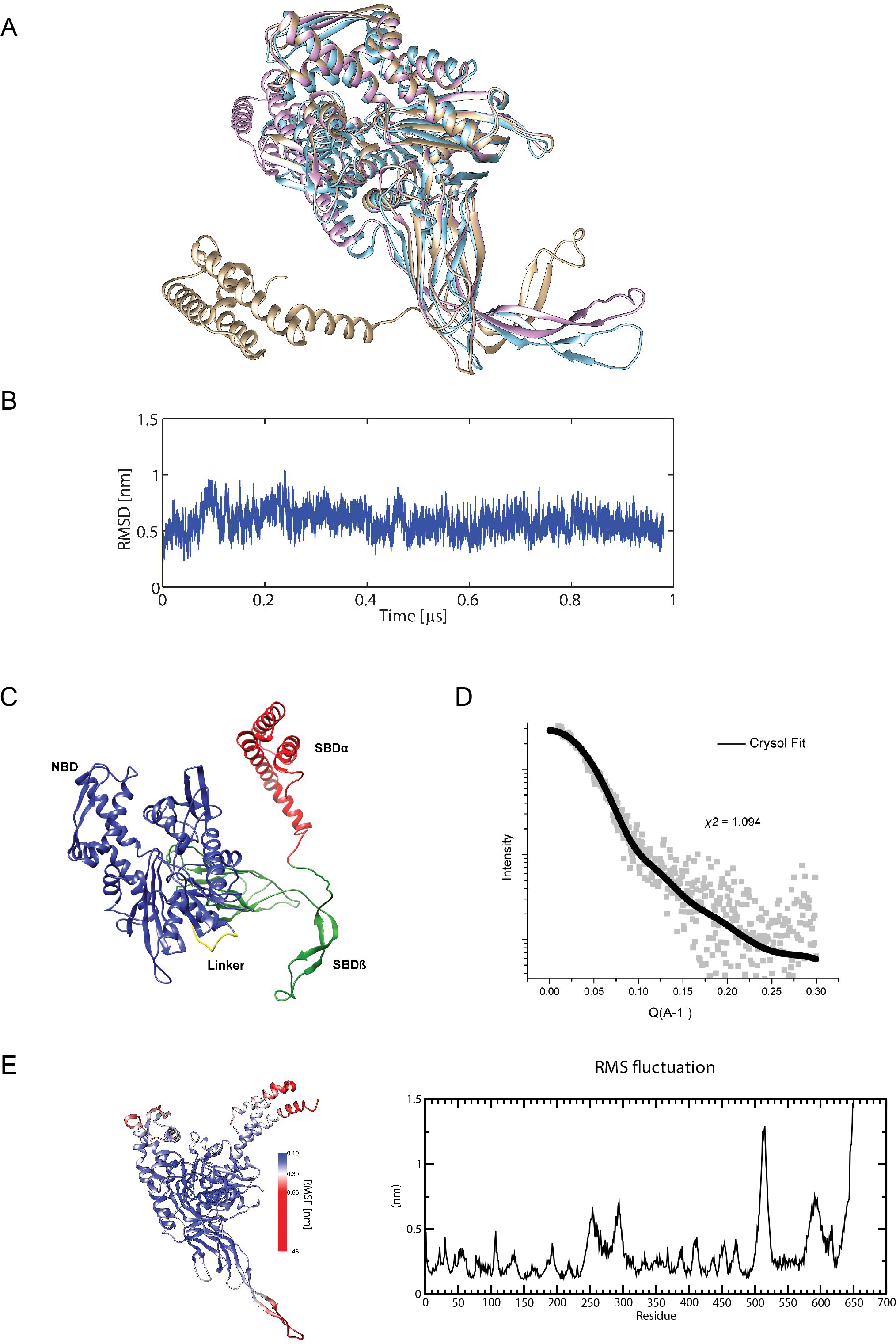
(**A**). Structural alignment of a) Sse1 crystal structure (PDB ID: 3C7N) (blue), b) molecular dynamics resolved final structure (pink) and SAXS-fitted structure (golden) reveals the maximum motion at the SBD domain. The SAXS-fitted model also shows an extension of the last ~110 residues at the C-terminal. (**B**) **Root mean square deviation (RMSD)**: Root mean square deviation was calculated for the all-atomistic 1 microsecond molecular dynamics simulation of native Sse1 and the structure was found stable post ~100ns. (**C**) Cartoon representation of MD stabilized final structure of Sse1 in Apo form. NBD has been depicted in blue, inter-domain linker in yellow, SBD-ß in green and SBD-α in red.(**D**) CRYSOL fitting of SREFLEX refined MD stabilized structure of Apo form of Sse1 with the experimental SAXS intensity profile gave *X*^2^ value of 1.094 suggesting a good fit with the experimental SAXS profile. (**E**). **Molecular dynamics simulation of native Sse1 reveals highly dynamic regions: Left panel:** Root mean square fluctuation of the protein reveals C-terminal helices and part of the beta-sheet region to be highly dynamical. **Right panel**: Residue-wise RMS fluctuation.

**Figure S2.**
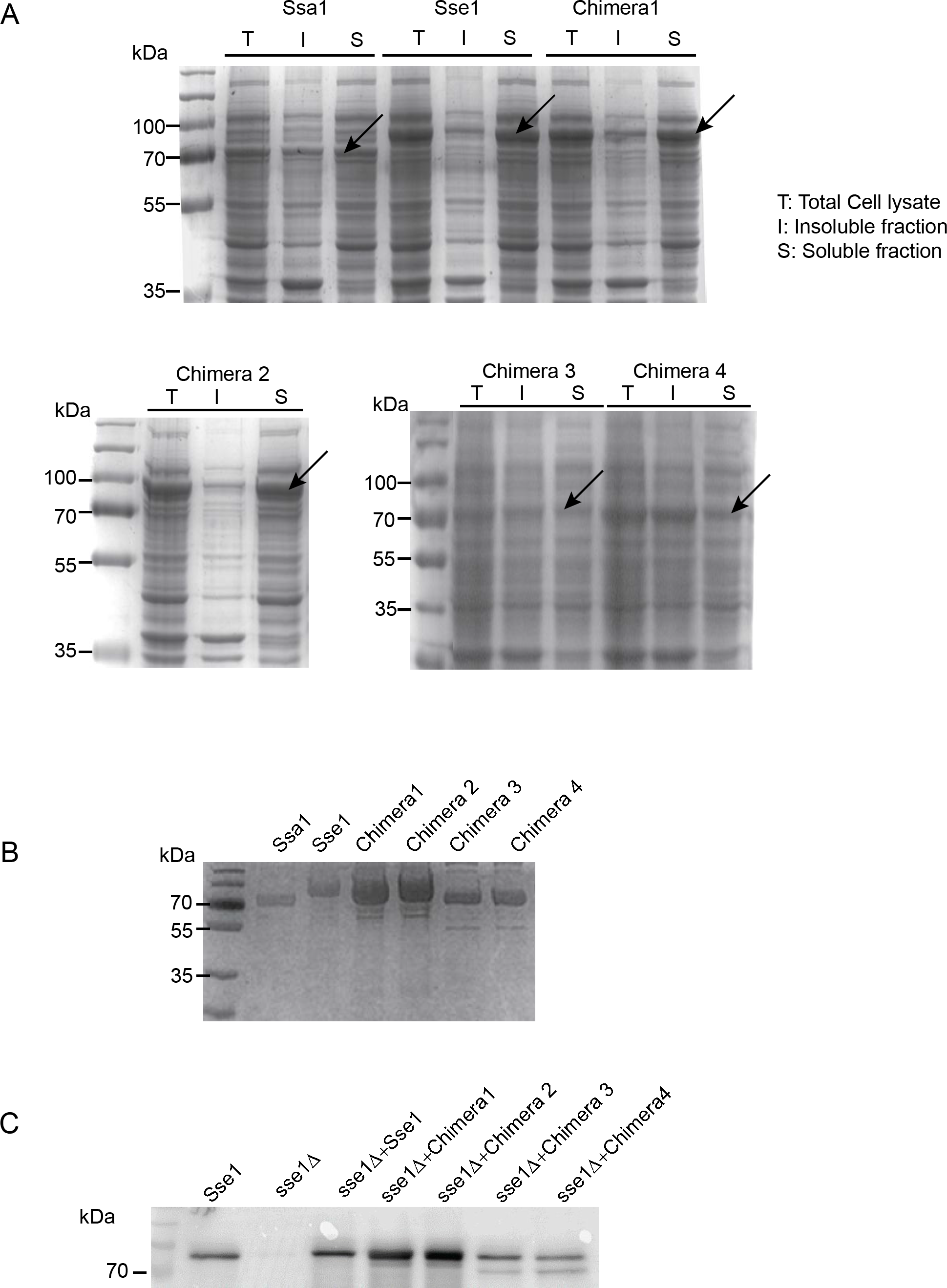
(A). Solubility assay for wild-type Ssa1, Sse1 and Hsp110-70 Chimeric proteins: The expression and the solubility of the wild-type Sse1 and all the chimeras were assayed by fractionating the whole cell lysate of BL21-DE3 cells into soluble (S) and insoluble fractions (I) by centrifugation. The total cell lysate before (T) and the soluble (S) and insoluble fractions (I) were run in SDS-PAGE and the gels were stained with Coomassie brilliant blue (CBB). The overexpressed protein bands have been shown with an arrow in the respective soluble fraction lanes. (B). The wild-type and chimeric proteins of Sse1 and Ssa1 were purified as described by in materials and methods section. The CBB stained SDS-PAGE shows the purified proteins. (C). The expression of wild-type Sse1 and all the chimera were checked by expressing all the proteins from overexpression vector (pJV340) in the *sse1*Δ cells and detected with anti Sse1 polyclonal antibody by western blot.

**Figure S3.**
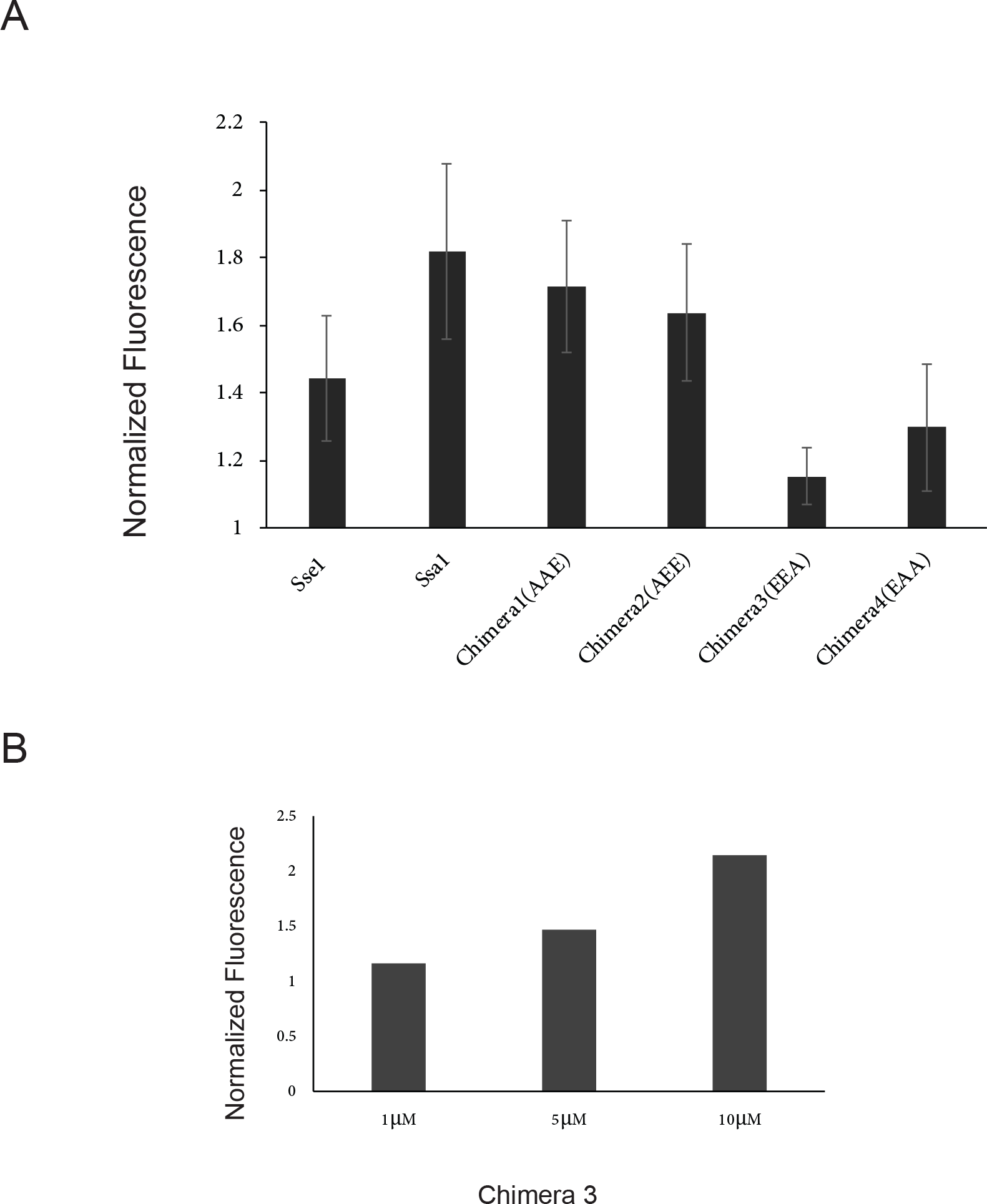
ATP binding assay for wild-type Ssa1, Sse1 and Hsp110-70 Chimeric proteins (A). The Nucleotide binding affinity for wild-type Ssa1, Sse1 and Hsp110-70 Chimeric proteins were assayed using mant ATP (2'-(or-3')-O-(N-Methylanthraniloyl) Adenosine 5'-Triphosphate). The ATP binding capacity of wild-type Sse1 and Chimeras were measured following the increase in fluorescence (Excitation/Emission: 355nm/428nm) after binding to the proteins by incubating 100nm of MANT-ATP (fluorescent analog of ATP) and 1μM of protein (wild-type Sse1, Ssa1 and Chimeras) in binding buffer containing 25mM HEPES, pH 7.4, 150mM KCl and 5mM MgCl_2_. The relative fluorescence was calculated as the ratio of fluorescence emission of mant-ATP bound to protein of interest and the emission of free mant-ATP. **(A).** shows that chimera 1 and 2 show ATP binding affinity is equivalent to Ssa1 (the NBD of Chimera 1 and 2 are obtained from Ssa1) and similarly, chimera4 shows similar binding affinity like Sse1 (the NBD of chimera 3 and 4 are obtained from Sse1). In contrast, chimera3 exhibited less affinity towards ATP. **(B).** The binding affinity of Chimera3 with mant-ATP with increasing concentrations of the proteins were checked and we found consistent increase of binding to mant- ATP with increasing concentrations of the protein showing less affinity of this chimera towards ATP.

